# Perceptual decisions about object shape bias visuomotor coordination during rapid interception movements

**DOI:** 10.1101/821074

**Authors:** Deborah A. Barany, Ana Gómez-Granados, Margaret Schrayer, Sarah A. Cutts, Tarkeshwar Singh

**Author notes:** Corresponding author: Tarkeshwar Singh, Department of Kinesiology, Behavioral and Cognitive Neuroscience Program, 330 River Rd, 115H, Athens, GA-30605, USA.

## Abstract

Visual processing in parietal areas of the dorsal stream facilitates sensorimotor transformations for rapid movement. This action-related visual processing is hypothesized to play a distinct functional role from the perception-related processing in the ventral stream. However, it is unclear how the two streams interact when perceptual identification is a prerequisite to executing an accurate movement. In the current study, we investigated how perceptual decision-making involving the ventral stream influences arm and eye movement strategies. Participants (*N* = 26) moved a robotic manipulandum using right whole-arm movements to rapidly reach a stationary object or intercept a moving object on an augmented-reality display. On some blocks of trials, participants needed to identify the shape of the object (circle or ellipse) as a cue to either hit the object (circle) or move to a pre-defined location away from the object (ellipse). We found that during perceptual decision-making, there was an increased urgency to act during interception movements relative to reaching, which was associated with more decision errors. Faster hand reaction times were correlated with a strategy to adjust the movement post-initiation, and this strategy was more prominent during interception. Saccadic reaction times were faster and initial gaze lags and gains greater during decisions, suggesting that eye movements adapt to perceptual demands for guiding limb movements. Together, our findings suggest that the integration of ventral stream information with visuomotor planning depends on imposed (or perceived) task demands.

**New and Noteworthy:** Visual processing for perception and for action are thought to be mediated by two specialized neural pathways. Using a visuomotor decision-making task, we show that participants differentially utilized online perceptual decision-making in reaching and interception, and that eye movements necessary for perception influenced motor decision strategies. These results provide evidence that task complexity modulates how pathways processing perception versus action information interact during the visual control of movement.

## Introduction

Many functional sensorimotor skills require rapid visual processing and perceptual decision-making. A very commonly encountered situation during driving is when drivers must decide whether to yield or stop at an intersection. The decision should be made from a distance by judging the shape of the sign at an intersection. If the shape is judged as a stop sign, the driver would slowly press their foot on the brake to bring the car to a gradual stop. However, if the shape is judged as a yield sign, the driver might just slow down or even hit the accelerator if there is no incoming traffic. The driver’s ability to make the correct decision and movement depends on efficient real-time processing of visual sensory information in the two visual processing streams (Goodale and Milner 1992; Mishkin et al. 1983). The distance between the sign and the car, the presence of other incoming traffic, and the associated motor actions are likely processed by the posterior parietal cortex along the dorsal visual stream (Culham et al. 2006; Rizzolatti et al. 2002; Rizzolatti and Matelli 2003). The shape and symbols on the sign are perceived by the lateral occipital and inferior temporal cortex along the ventral visual stream (Ales et al. 2013; Grill-Spector et al. 2001; Lehky and Tanaka 2016; Schwartz et al. 1983). Though the contributions of these streams to visuomotor and visuoperceptual processing is well delineated, it is still unclear how these two streams interact and process sensory information in real-time to facilitate rapid visuomotor actions.

The goal of the present study was to understand how engaging the ventral stream affects the spatiotemporal course of movement selection and execution. Many behavioral (reviewed in Gallivan et al. 2018; Hecht et al. 2008; Rosenbaum et al. 2007; Song and Nakayama 2009) as well as neurophysiological studies (reviewed in Cisek and Kalaska 2010) have provided empirical support for simultaneous specification of competing motor plans in the dorsal visual stream. Rapid movement modifications have also been shown when a perceptual decision is made based on ventral stream related attributes, such as object color or shape (Cressman et al. 2007; Schmidt 2002; Song and Nakayama 2008; Veerman et al. 2008), though at slower time scales than motor decisions based on dorsal stream processing of spatial or motion-related properties (Day and Lyon 2000; Franklin et al. 2016; Gritsenko et al. 2009; Sarlegna and Mutha 2015). These results imply that despite functionally segregated roles, goal-directed visuomotor actions ultimately necessitate online interaction between the ventral and dorsal streams (Gallivan and Goodale 2018; Milner 2017; Song and Nakayama 2009).

In these previous studies, the movement required is typically a simple reach executed to a spatially defined goal. However, the capacity for integration of ventral stream information with online decision-making and motor planning may depend on the computational complexity of the movement (van Polanen and Davare 2015). In contrast to simple reaching, interception movements present a challenge for the motor system due to the uncertainty in estimating the velocity and future position of the target and in specifying an appropriate motor plan to hit the target at the desired time and location (Brenner and Smeets 2009; Merchant et al. 2009; Zago et al. 2009). Humans can achieve high interception accuracy via continuous updating of movement trajectories under visual feedback control (Brenner and Smeets 2018), but it is unclear how these interception mechanisms may be modulated by perceptual decision processes mediated by the ventral visual stream.

In the present study, we developed a rapid visuomotor decision-making task where participants were asked to make reaching or interception movements under relatively fast or slow time constraints. In some blocks of trials, participants were simply required to hit a stationary (reaching) or moving (interception) object as quickly and as accurately as possible. In separate blocks, participants needed to select among two alternative actions that required correctly identifying the object’s shape (hit the circle and avoid the ellipse). Our first hypothesis was that engaging the ventral stream would elicit stronger interference between ventral and dorsal stream processes during interception than reaching movements. We predicted that both decisional and aiming accuracy would be lower for interception movements.

In contrast to fixations on static targets during reaching movements, smooth-pursuit eye movements track moving targets and engage additional neural resources (Lencer and Trillenberg 2008; Lisberger 2015) during interception movements. Once the moving target is stabilized on the retina, the limb motor system may rely on oculomotor efferent signals during pursuit eye movements to perform continuous retinotopic to limb-centric coordinate transformations (Gauthier et al. 1990) and guide limb movements. The neural regions involved in eye movement processing overlap with those involved in decision-related signals (Fooken and Spering 2019; Gold and Shadlen 2007; Heekeren et al. 2008; Joo et al. 2016), and this likely affects recognition of object features during fast smooth-pursuits (Ludvigh and Miller 1958a; Schutz et al. 2009; Westheimer and McKee 1975). Thus, our second hypothesis was that when the ventral stream is engaged during interception movements, the oculomotor signature of pursuit eye movements will change. Specifically, we expected higher gaze gains (computed as ratio of gaze velocity and target velocity) during perceptual decisions.

## Methods

### Participants

Twenty-six healthy, right-handed participants (16 women; 23.7 ± 5.5 years) completed the experiment. All participants had no known history of neurological disorders and had normal or corrected-to-normal vision. Each participant provided written informed consent prior to participating and were compensated for their participation. All study procedures were approved by the Institutional Review Board at the University of Georgia.

### Apparatus

Participants were seated in a chair and used their right hand to grasp the handle of a robotic manipulandum that could move in a horizontal plane (KINARM End-Point Lab, BKIN Technologies, Kingston, Ontario, Canada) (see Fig.1A). All visual stimuli were projected at 60 Hz onto a semi-transparent mirror from a monitor above the workspace. This set-up allowed the stimuli to appear on the same horizontal plane as the handle and to occlude direct vision of the hand. During task performance, the robot applied a small background load (−3 N in the Y direction) to the handle and recorded movement position and velocity at 1000 Hz. The monocular eye position of each participant was recorded at 500 Hz using a video-based remote eye-tracking system (Eyelink 1000; SR Research, Ottawa, ON Canada) integrated with the robot and calibrated for the 2D horizontal workspace. Data from the eye-tracker and robot were time-synced offline using MATLAB (version 9.5.0; The MathWorks, Natick, MA).

**Figure 1:**
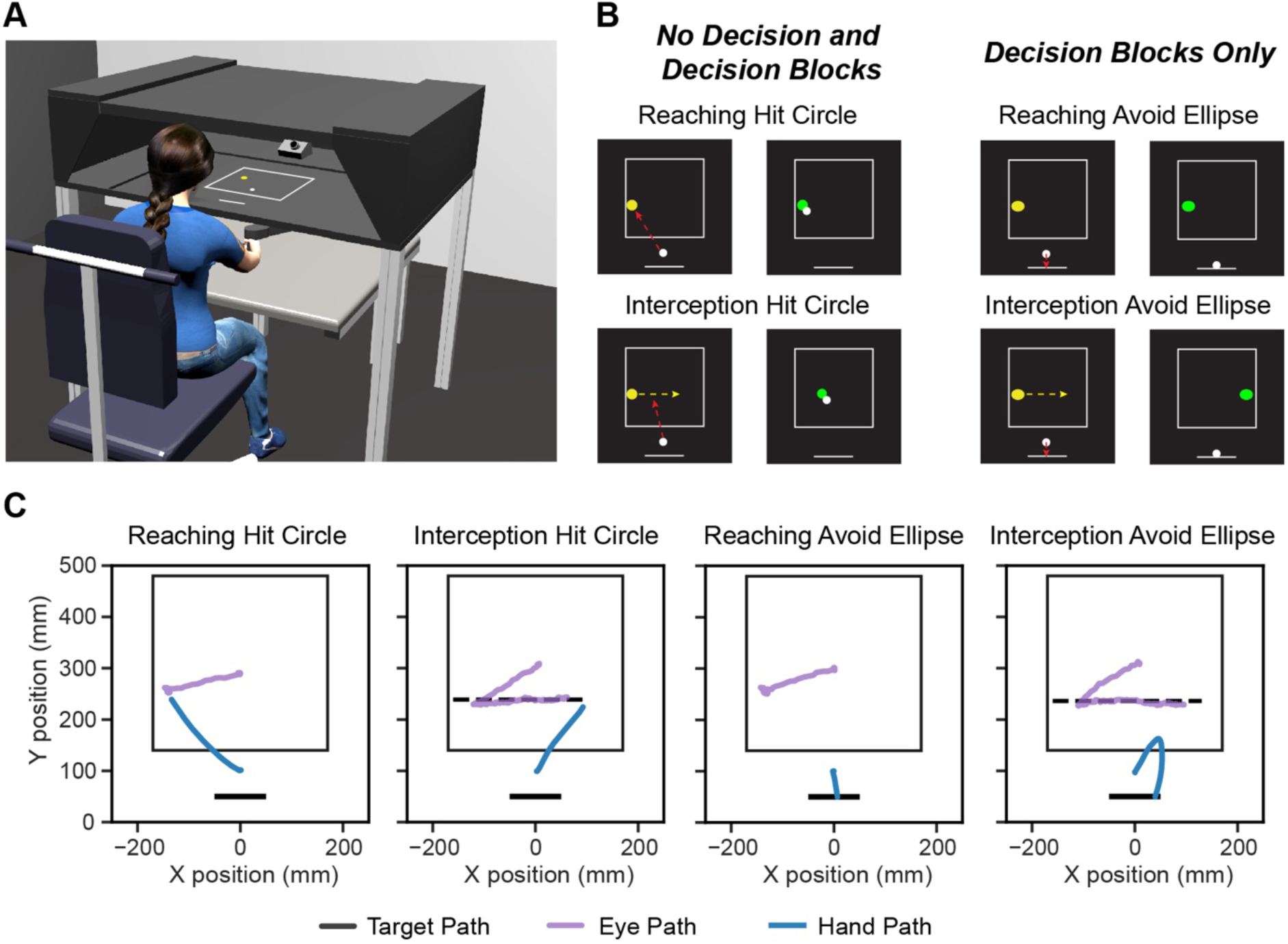
Experimental design and example trials. A: Experimental setup. Participants moved a robotic manipulandum with their right hand to control a cursor (white circle) in response to an object (yellow circle) on the visual display. A remote gaze-tracker at the back of the workspace recorded eye positions in Cartesian coordinates of the workspace. B: Trial types. On every trial, participants were instructed to hit or avoid depending on object shape (hit circle, avoid ellipse). No Decision blocks consisted of only circles; Decision blocks mixed circle and ellipse trials with equal probability. Participants either reached a stationary object (Reaching) or intercepted a moving object (Interception). The object turned green for correct hits (circle hits) and red for incorrect (if ellipses were hit). Similarly, if a circle was missed, it turned red at the end of the trial (Fast blocks trial duration: 800 ms; Slow blocks: 950 ms), and if movement was made towards the bar when an ellipse appeared in the workspace, it turned green at the end of the trial. C: Sample 2D eye and hand paths for each trial type from a representative participant.

### Experimental design and procedure

Participants performed rapid whole-arm reaching and interception movements in which they were instructed to either hit or avoid an object based on the object’s shape. At the beginning of each trial, participants moved a cursor (white circle, 1 cm diameter) representing their veridical hand position to a start position (yellow circle, 2 cm diameter) located at the midline of the visual display (x=0). After reaching the start position, a fixation cross appeared at the midline 22 cm from the start position in the Y direction. Participants were required to maintain fixation and keep their hand at the start position for 500 ms, after which the fixation cross and start position disappeared.

Following a fixed 200 ms delay, a yellow object was presented on the display near either the left or right edge of a rectangular box (34 × 34 cm) centered on the midline and 22 cm above the start position (see Fig. 1B). The possible object shape on a given trial, and the participant’s task, depended on the experimental block. During No Decision blocks, participants were informed that the object shape would always be a circle (2 cm diameter), and that they should hit the circle as quickly and as accurately as possible. A “hit” was recorded when the cursor first touched the circle—participants were not required to stop at the circle. During Decision blocks, participants were informed that the object would appear as either a circle or an ellipse (major axis = 2.3 cm; minor axis = 2 cm) with equal probability. The lengths of the ellipse axes were selected based on pilot experiments to ensure that the object must be foveated to differentiate it from a circle. As in the No Decision blocks, if the participants saw a circle, they were instructed to hit it as quickly and as accurately as possible. However, if an ellipse appeared, participants were instructed to avoid hitting the ellipse and instead move in the opposite direction toward a horizontal bar (10 cm width) centered on the midline and −4 cm from the start position in the y direction (see Fig. 1B). Thus, in contrast to No Decision blocks, in which participants could simply plan to hit the object on every trial, Decision block trials required the participant to accurately identify the object shape in order to perform the correct action (i.e., hit the circle or avoid the ellipse). Therefore, in addition to the No Decision blocks, the Decision condition required two additional steps, object identification and selection of an appropriate motor plan.

For each block of trials, the object either moved horizontally across the display (Interception) or remained in the same position (Reaching). On Interception trials, the object appeared ±16 cm to the left or right of the midline (Y position range 14.5 - 17 cm from the start position, uniform distribution) and traversed at a constant Euclidean velocity of ±40 cm/s (Fast) or ±34 cm/s (Slow) toward the other horizontal boundary of the rectangular box. The varying object velocity was added to test the hypotheses under stricter conditions of time constraints. On Reaching trials, the object appeared to the left or right of the midline with starting positions drawn from a uniform distribution (X position range: ±13 - 16 cm from midline; Y position range 14.5 - 17 cm in front of start position) and remained stationary. For both types of trials, the object remained on the visual display until it was hit or for the maximum trial duration. On Interception trials, the maximum trial duration equaled the time it took for the object to arrive at the horizontal boundary given its velocity: 800 ms for fast velocities (±40 cm/s) and 950 ms for slow velocities (±34 cm/s). To match the Interception trial durations, objects remained on the screen for a maximum of 800 ms (Fast) or 950 ms (Slow) during Reaching trials. Before each block, participants were informed about the object motion (moving or stationary) but were not given any information about the object speed or trial duration.

Performance feedback was provided for 500 ms once the object was hit (i.e., the cursor overlapped with the object) or the maximum trial duration was reached. If a circle was correctly hit, the circle would turn green; if the circle was missed it would turn red. An ellipse would turn red if it was incorrectly hit instead of avoided and would turn green if correctly avoided. The next trial began following a 1500 - 2000 ms delay.

Participants performed 8 experimental blocks of 90 trials each (720 trials total). Block order was counterbalanced across participants. Each experimental block consisted of a unique combination of decision type (No Decision or Decision), movement type (Reach or Intercept), and trial duration (Fast or Slow). Object shape (during Decision blocks) and the object start location were randomized across trials within each block.

### Data Analysis

All hand and eye movement data were analyzed using MATLAB (version 9.5.0, The MathWorks, Natick, MA) and Python (version 3.7). Statistical analyses were performed in R (version 3.6.0).

### Arm Movements

Hand position and velocity data were first smoothed using a fourth-order Butterworth low-pass filter with a 5 Hz cutoff. Movement onset was defined as the time the tangential velocity first exceeded 5% of the first local peak. Reaction time (RT) was calculated as the time from appearance of object in the workspace to movement onset. Trials were excluded if there was no identifiable RT or if RT was less than 100 ms (1.4% of all trials). Trials were also excluded if participants received correct feedback despite inaccurate motor performance; this was the case when the participant hit the circle only after missing the object on the initial attempt (2.3% of all trials). Peak speed (PS) was defined as the maximum tangential velocity of the hand position at the first local peak. Since PS could differ depending on the object decision in Decision blocks, only trials in which the participant continually moved toward the circle throughout the trial were included (49.3% of all Decision trials).

For each trial, we examined the hand kinematics to determine decisional and motor performance accuracy at different stages of the movement. The initial direction (ID) of the movement was calculated as the angle between the midline and the vector linking the hand position at the start to the hand position at peak acceleration. In Decision blocks, the initial decision was based on the ID of the movement: movements were classified either as being aimed toward the object or toward the bar. Initial decision errors were computed for each participant as the percentage of trials in which the initial decision did not match the expected movement direction given the true object identify (i.e., aimed toward the bar on trials with a circle or aimed toward the object on trials with an ellipse). Likewise, final decision errors were calculated as the percentage of trials the participants’ final hand position was closer to the bar on circle trials or closer to the object on ellipse trials. Trials in which the initial decision and the final decision were different (e.g., aimed toward the circle but attempted to hit the bar) were classified as “redirect” movements, indicating a change-of-mind after movement initiation (Resulaj et al. 2009). We quantified both the total percentage of redirect movements across all Decision trials, as well as the percentage of initial decision errors that were redirected. This latter index characterizes how well participants were able to correct wrong initial decisions online.

Finally, to compare motor performance across No Decision and Decision blocks, we calculated aiming accuracy on trials continually directed toward the circle (i.e., all valid No Decision trials and Decision circle trials in which both the initial and final decision were correct). An *aiming error* was defined as whenever the hand position reached the Y-position of the object, but nevertheless did not successfully hit the object before the trial elapsed.

### Eye Movements

Details of gaze processing and gaze-event identification are provided in more detail in previous work (Singh et al. 2017; Singh et al. 2016). Briefly, gaze data were low-pass filtered at 20 Hz and preprocessed to remove blinks, one-sample spikes (due to incorrect detection of corneal reflection), and screen outliers (due to instances when gaze drifts outside the workspace). Gaze events were identified as saccades and fixations using adaptive velocity and acceleration thresholds (Singh et al. 2016). Our previous analyses showed that velocity thresholds vary substantially between participants but that acceleration threshold is relatively constant (6,000°/s^2^). For each velocity peak that exceeded the velocity threshold, we confirmed that the peak acceleration leading up to the velocity peak also exceeded the acceleration threshold. If both thresholds were exceeded, we classified the gaze event as a saccade. For each saccade, we found the first inflection point before and after the local peak in gaze angular velocity. Saccade onset corresponded to the first inflection point before the local peak in gaze angular velocity. Saccade offset was determined by starting at the first inflection point after the local peak in gaze angular velocity and finding the first point in time at which the gaze velocity and acceleration remained continuously lower than the respective thresholds for at least 40 ms.

For interception movements, smooth-pursuits were identified when gaze and target locations and velocities were continuously within a *foveal visual radius* as described in Singh et al. (2016). Briefly, because targets were presented in a transverse plane, the foveal visual radius accounts for larger spatial distances for the same foveal visual acuity (2-3°) when the objects were presented farther away from the body. Note that a gaze event was only classified as a smooth-pursuit if the target was foveated. Individual saccades were discarded if the duration was <5 ms, and smooth-pursuits/fixations were discarded if the duration was <40 ms. On some trials, participants made predictive saccades anticipating the location of the object. Since we were only concerned with visually-guided performance, we eliminated any saccade initiated <100 ms after target onset and any initial saccade not directed to the object (>100 mm from object). Following exclusion of individual saccades, we defined a valid trial for the task as one containing an initial saccade to the target followed by a fixation or smooth-pursuit. Thus, gaze for a trial was not analyzed if the trial did not contain a valid saccade and a gaze event (fixation or pursuit) or if a gaze event (fixation or pursuit) occurred before any saccade. Overall, gaze data were included for 90.7% of Reaching trials and 88.6% of Interception trials. Data from two subjects were not included in the eye movement analyses because fewer than 50% of their trials were identified as valid according to the above criteria.

Saccadic reaction time (SRT) for both Reaching and Interception trials was calculated as the onset of the initial saccade for a given trial. For interception movements, we also determined the gaze lag as the horizontal distance (mm) between the moving object and the eye position at the end of the first saccade, and throughout the gaze duration (excluding catch-up saccades occurring during the smooth-pursuit period). Gaze gain was calculated as the gaze angular velocity divided by the object angular velocity and average gain was quantified for the open-loop (15-100 ms of gaze), first 100 ms of the closed-loop (next 100 ms of gaze), and full closed-loop (gaze after first 100 ms) phases (excluding catch-up saccades). Gaze gain for the first 15 ms was not analyzed due to the potential for artificially high velocities from the offset of the preceding saccade. Removal of the first 15 ms did not affect differences in gaze gain across conditions. Of note, smooth-pursuit gains are typically computed using eye-trackers with chin rests (Brostek et al. 2017; Churchland and Lisberger 2002) or eye-trackers that are head-mounted (Spering et al. 2005). With these eye-trackers, gaze movements are computed as eye-in-head movements. In contrast, we used a remote eye-tracker which allowed small head movements to occur. Thus, we chose to report gaze gains instead of smooth-pursuit gains (Barnes 1993; Ranalli and Sharpe 1988). Finally, we determined the number of catch-up saccades as a function of time after gaze onset and quantified the average number of catch-up saccades during the entire gaze duration.

### Statistical Analyses

To assess how the introduction of perceptual decision-making influenced RT, PS, and SRT, we computed the means for each combination of decision type, movement type, object velocity, and object start location (left or right). We then subtracted the No Decision block means from the Decision block means, separately for each participant and movement type/trial duration combination. A one-sample *t*-test was used to determine whether the change between Decision and No Decision means were significantly different from zero, and a 2 (Reaching or Interception) x 2 (Fast or Slow) repeated-measures ANOVA assessed whether the effect of decision-making differed across movement type and trial duration. Measures of decision-making and hand and eye motor performance were assessed across conditions using repeated-measures ANOVAs. For all ANOVA tests, the alpha level was set at 0.05 and effect sizes are reported using generalized *η*^2^. Post hoc pairwise comparisons were conducted using the Holm correction (Holm 1979). Linear regression was used for bivariate comparisons, with alpha set to 0.05, and the statistical comparison of correlations between conditions was evaluated using the Dunn and Clark’s *z* for dependent groups with nonoverlapping variables (Dunn and Clark 1969), as implemented in *cocor* package in R (Diedenhofen and Musch 2015).

## Results

### Final decision errors occurred more frequently for interception than reaching movements

In the task, participants made rapid eye and arm movements in response to an object appearing on the visual display. As illustrated in Figure 1C, after object onset participants typically made saccades directly to the object, followed by fixation on a stationary object near the right or left edge of the display boundary (Reaching trials) or pursuit of an object moving at a constant Euclidean velocity from one boundary to the other (Interception trials). Participants either attempted to hit any circle that appeared by moving the cursor (representing hand position) to the object before the end of the trial or avoid any ellipse that appeared by moving in the opposite direction toward a bar on the display.

Figure 2A shows the hand trajectories for a representative participant. Each line indicates the hand path from object onset until the participant hit their intended target (object or bar), or until the maximum trial duration (if neither the object nor the bar was hit). During No Decision blocks, the object was always a circle, whereas in Decision blocks, the object could be either a circle or ellipse. The addition of the decision-making task component led to clear differences in where participants chose to intercept the object. In No Decision blocks, on average, participants tended to intercept the object slightly after it crossed the midline (*M =* 20.1 ± 5.9 mm from midline). In contrast, there was a significant shift in object hit locations during Decision blocks (*M =* 75.0 ± 5.6 mm from midline) [main effect of decision: *F*(1,25) = 228.77, *p* < 0.001, η^2^ = 0.66]. As expected, interceptions were made later when the object was moving faster [main effect of trial duration: *F*(1,25) = 110.32, *p* < 0.001, η^2^ = 0.13].

**Figure 2:**
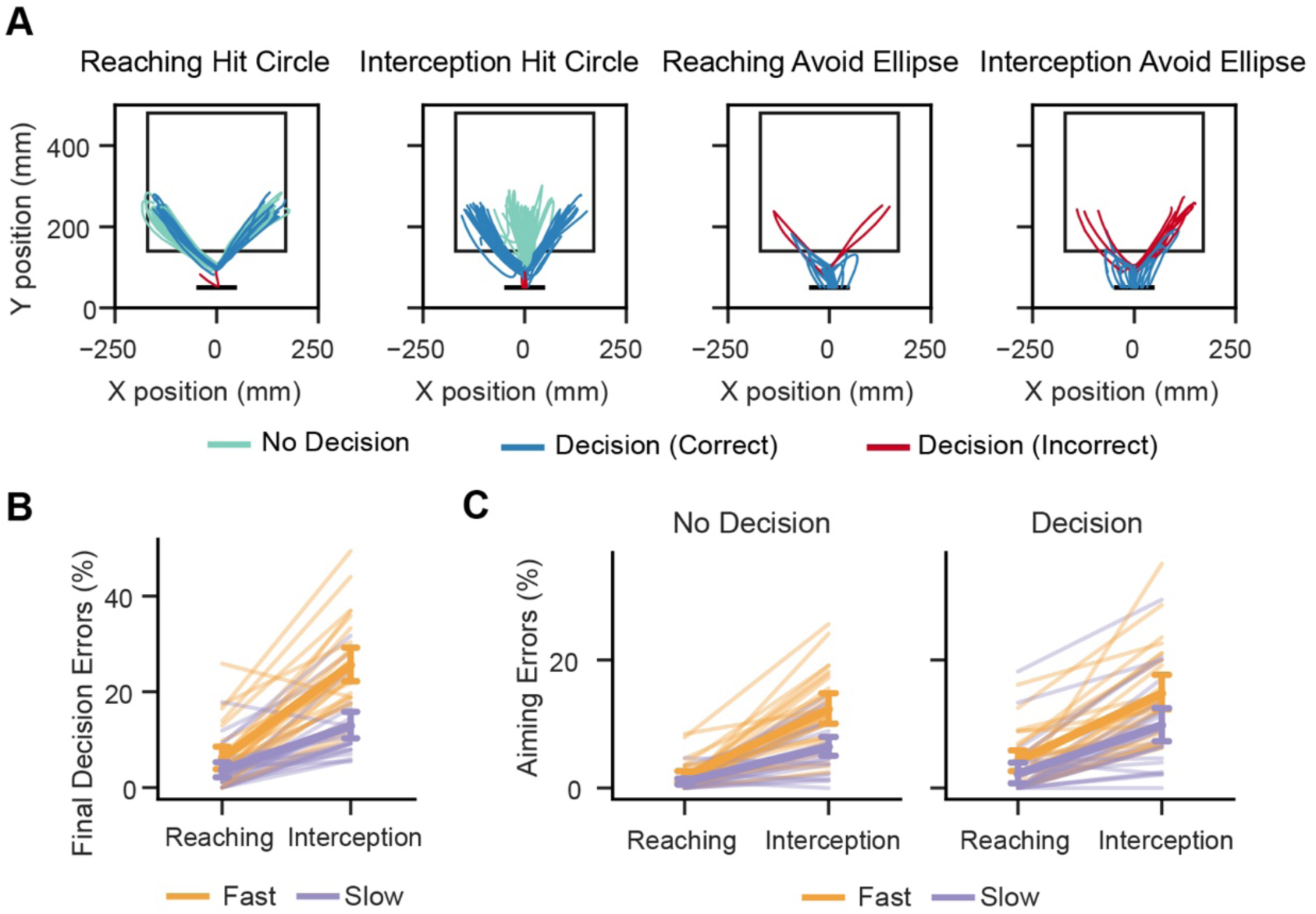
Final decision errors for interception and reaching movements. A: Sample hand paths from a representative participant. During No Decision blocks, participants were required to reach or intercept a circle appearing on the display (green paths, left two panels). During Decision blocks, participants were required to hit a circle if it appeared (blue and red paths, left two panels), or avoid an ellipse (right two panels). Final decisions on these trials were classified as correct if the final hand position was closer to the correct location (object or bar) given the object’s identify (blue paths), and incorrect if not (red paths). B: Final decision errors were higher for interception than reaching and for fast (800 ms) than slow (950 ms) trial durations. C: Aiming errors were higher for during interception, and aiming errors increased similarly for both reaching and interception during Decision blocks. Errors were calculated as the percentage of all trials in which the y-position of the object was reached but the object was not hit. Individual lines represent the means for one participant. Error bars show the 95% confidence interval of the group mean estimate.

In Decision blocks, final decisions were classified as either correctly attempting to hit the circle or avoid the ellipse, or incorrectly attempting to hit the ellipse or avoid the circle (Fig. 2B). The percentage of final decision errors was higher for interceptions than for reaching movements [main effect of movement type: *F*(1,25) = 113.03, *p* < 0.001, η^2^ = 0.52] and for faster trial durations [main effect of trial duration: *F*(1,25) = 107.72, *p* < 0.001, η^2^ = 0.23]. The increase in errors at faster durations was larger for interceptions [interaction of movement type and trial duration: *F*(1,25) = 47.38, *p* < 0.001, η^2^ = 0.12], indicating that faster object velocity reduced interception decision accuracy beyond decreasing the time possible to hit the object.

For Decision blocks, we then computed aiming errors for only those trials where the final decision was correct. As expected, the additional computational costs associated with estimating object velocity and movement timing led to more aiming errors during interception movements. In both No Decision and Decision blocks, there were a higher percentage of aiming errors for Interception [main effect of movement type: *F*(1,25) = 129.22, *p* < 0.001, η^2^ = 0.43], especially at faster trial durations [interaction of movement type and trial duration: *F*(1,25) = 20.88, *p* < 0.001, η^2^ = 0.04], reflective of the greater difficulty in intercepting an object at higher speeds (Fig. 2C). There was an increase in aiming errors in Decision blocks [main effect of decision: *F*(1,25) = 11.49, *p* = 0.002, η^2^ = 0.06], but the increase did not differ between Reaching and Interception [interaction of movement type and decision: *F*(1,25) = 1.48, *p* = 0.24, η^2^ = 0.003]. Together, these results suggest that during time-constrained perceptual decision-making, the added task demands of interceptive movements affected the decisional accuracy more than the motor accuracy.

### Perceptual decisions increase urgency to act more for interception relative to reaching

One potential strategy participants could have employed in the Decision trials is to complete the recognition of the object shape before initiating a movement. Such a strategy would minimize an erroneous commitment to a movement that would later have to be reversed. If this were the case, initial decisions should have been similar between Reaching and Interception movements. In contrast, there was a large increase in initial decision errors during Interception relative to Reaching [main effect of movement type: *F*(1,25) = 121.09, *p* < 0.001, η^2^ = 0.48] (Fig. 3A). Most of these errors (91.4 %) were due to initially aiming toward the ellipse (which had to be avoided), suggesting a default initial strategy of trying to hit rather than avoid the object and then correct the movement if the object shape was correctly identified during the movement. This default strategy was used more often during faster trials [main effect of trial duration: *F*(1,25) = 19.09, *p* < 0.001, η^2^ = 0.05], when there were greater constraints to hit the object in time.

**Figure 3:**
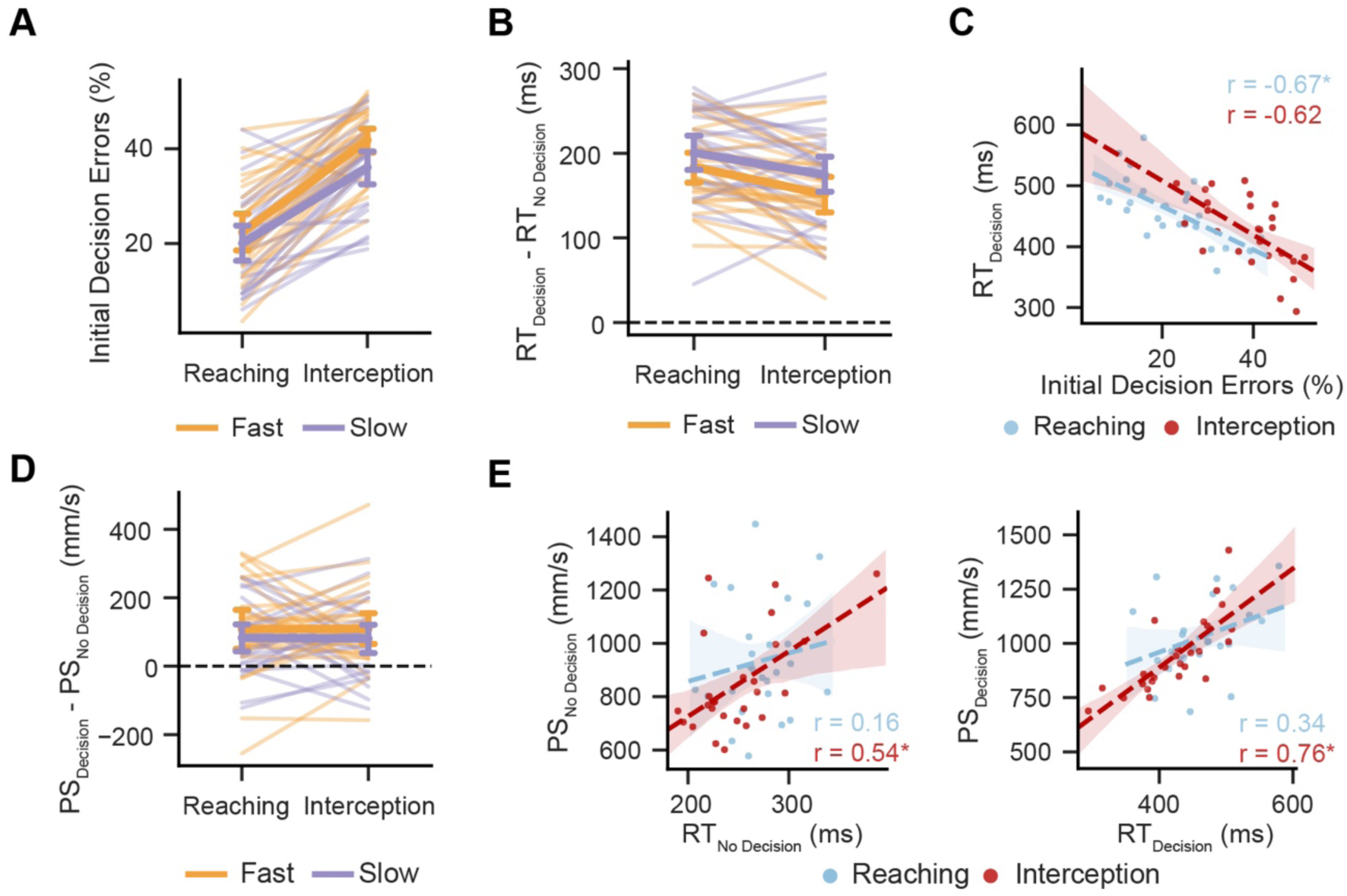
Reaction times and limb kinematics for interception and reaching movements. A: Initial decision errors were higher for interception and for fast (800 ms) trial durations. B: The increase in reaction time from No Decision to Decision blocks was smaller for interception relative to reaching. C: Participants were shorter reaction times during Decision blocks were exhibited a higher number of initial decision errors. D: Peak speed increased for Decision blocks similarly for reaching and interception. E: Reaction time and peak speed tended to be more correlated during interception and Decision blocks. For line plots, individual lines represent the means for one participant and error bars show the 95% confidence interval of the group mean estimate. For regression plots, each dot represents the mean value for one participant and shaded area represents the 95% confidence interval of the regression estimate. * indicates p < 0.05.

As expected, perceptual decision-making led to a significant reaction time (RT) delay. Relative to No Decision blocks, RTs for Decision blocks were on average 178 ± 11 ms longer [*t*(1,25) = 20.04, *p* < 0.001] (Fig. 3B). Thus, perceptual decisions based on ventral stream processing clearly increased the time taken for object identification (circle or ellipse) and motor response selection (hit or avoid). However, the increase in RT for the Decision blocks differed depending on the type of movement and time constraints: RT increase was smaller for Interception [main effect of movement type: *F*(1,25) = 13.63, *p* = 0.001, η^2^ = 0.07], and for Fast movement blocks [main effect of trial duration: *F*(1,25) = 9.83, *p* = 0.004, η^2^ = 0.04]. This suggests that even though decisions added processing time, participants chose to limit pre-movement processing time when an interception was required or under more restrictive time constraints. The increased urgency to act came at the expense of initial decision accuracy: participants with shorter RTs during Decision blocks exhibited more initial decision errors for both reaching and interception movements (Reaching: *r* = −0.67, *p* < 0.001; Interception: *r* = - 0.62, *p* <0.001) (Fig. 3C).

During decision-making, there was also an increase in the speed of the response: on average, peak speed (PS) of movements attempting to hit the object increased by 95.4 mm/s [*t*(1,25) = 5.46, *p* < 0.001] (Fig. 3D). The change in PS did not vary based on movement type [main effect of movement type: *F*(1,25) = 0.00, *p* = 0.98, η^2^ < 0.01] or trial duration [main effect of trial duration: *F*(1,25) = 2.70, *p* = 0.11, η^2^ = 0.01]. For reaching movements, the increase in PS may reflect a general urgency to complete the movement more quickly after a prolonged decision period. For interception movements, where participants have a salient visual cue for time remaining (the object approaching the boundary), changes in PS are likely more directly related to changes in RT: the longer the participant waited to initiate movement, the less time available and longer movement amplitude necessary to hit the object. Indeed, for both No Decision and Decision blocks, there was a significant positive correlation between PS and RT (No Decision: *r* = 0.54, *p* = 0.003; Decision: *r* = 0.76, *p* < 0.001), which was not the case for reaching movements (No Decision: *r* = 0.16, *p* = 0.43; Decision: *r* = 0.34, *p* = 0.08) (Fig. 3E). The PS-RT correlation was significantly greater for Decision, Interception blocks than for No Decision, Reaching blocks (*z* = 2.98, *p* = 0.003), indicating that the lower RTs during decision-making for interception may be in part to allow for slower, shorter movement trajectories. Overall, the results suggest that perceived time constraints—amplified during both interception movements and faster trial durations—encourage earlier movement initiation even if the decision process is incomplete.

### Interception strategies favor ongoing decision-making after movement initiation

To further investigate how movements are planned relative to time-sensitive decision processing, we analyzed how often participants adjusted their movements online. To do this, we distinguished between “direct” and “redirect” movements. Direct movements were when both the initial and final decisions were directed toward the object (direct object hit) or to the bar (direct avoid). Redirect movements occurred when the final decision differed from the initial decision: as can be seen in Figure 2A, redirects were predominantly observed when the participant made an initial decision toward the object, only to curve back around to hit the bar (redirect-to-avoid). The opposite pattern—moving to the object after initially moving to avoid it (redirect-to-hit), rarely occurred (<0.01% of Decision trials), highlighting the greater accuracy demands imposed by hitting the object vs. hitting the bar.

All participants had both direct and redirect movements, indicating a mixture of strategies used during the task. Overall, redirect movements were more common during Interception [main effect of movement type: *F*(1,25) = 16.82, *p* < 0.001, η^2^ = 0.11], especially at Slow trial durations [interaction of movement type and trial duration: *F*(1,25) = 9.61, *p* = 0.005, η^2^ = 0.03] (Fig. 4A). This suggests that decisions about object shape could be modified after movement initiation. Furthermore, participants were more likely to rely on this strategy for complex interceptive movements and when there was more time for online corrections (Slow trials).

**Figure 4:**
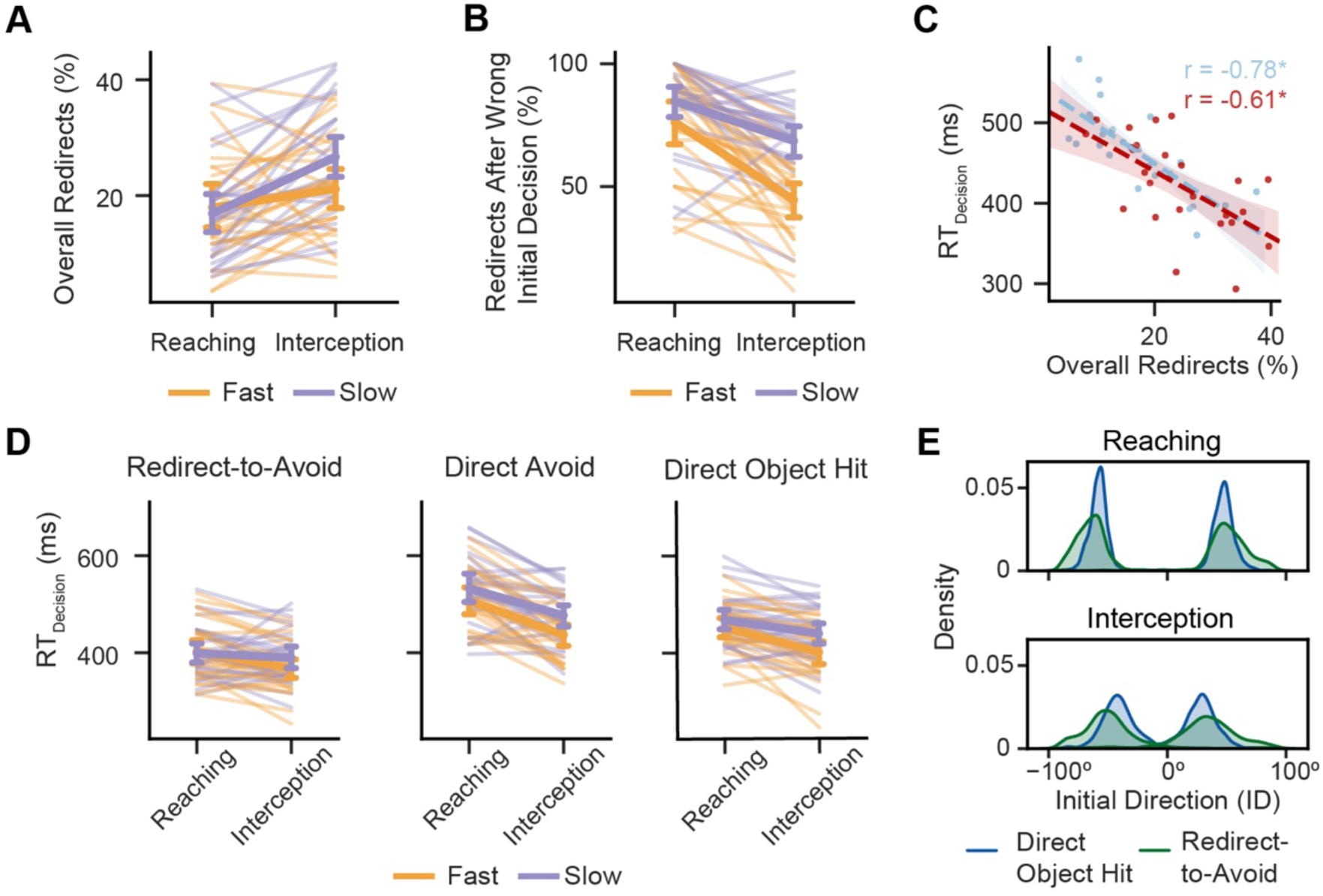
Redirected movements and Initial Directions (ID) reveal ongoing decision-making after movement initiation. A: Redirect movements (change between initial and final decision) during Decision blocks were higher for interception, suggesting more online adjustments after movement initiation. B: Initial decision errors were more likely to be corrected for reaching and slow trial durations. C: Participants were shorter reaction times during Decision blocks were exhibited a higher number of redirect movements. Each dot represents the mean value for one participant and shaded area represents the 95% confidence interval of the regression estimate. * indicates p < 0.05. D: Mean reaction times were shortest for redirect-to-avoid movements (initially aimed toward object then redirected to bar), longest for direct avoid movements (directed toward bar throughout), and intermediate for direct object hits (directed to object throughout). In all cases, interception reaction times were shorter than those for reaching. Individual lines represent the means for one participant and error bars show the 95% confidence interval of the group mean estimate. E: Kernel density estimate of the initial movement direction (0° = aimed at midline) for redirect-to-avoid and direct object hit movements. IDs were aimed farther from the midline for redirect-to-avoids during Decision blocks for both reaching (upper panel) and interception (lower panel).

Though redirect movements were used more during Interception, they were employed more effectively during Reaching. As shown in Figure 4B, after an initial decision error, a correct redirect of an initially wrong decision was more likely to occur for Reaching [main effect of movement type: *F*(1,25) = 50.82, *p* < 0.001, η^2^ = 0.30] and for Slow trial durations [main effect of trial duration: *F*(1,25) = 55.83, *p* < 0.001, η^2^ = 0.16]. Therefore, task difficulty limited the ability to implement a corrective movement when they were necessary.

If initial decisions were less likely to be corrected, why were participants more likely to redirect their movements during Interception trials? In Decision blocks, movements might have been initiated early (during both Reaching and Interception trials) before the perceptual decision was complete, but once the movements were underway the complexity of the interception movements may have made it much harder to correct them. If this is the case, initiation of redirect movements should be associated with shorter RTs. Indeed, for both Reaching and Interception, participants with a higher proportion of redirect movements exhibited shorter decision RTs [Reaching: *r* = −0.78, *p* < 0.001; Interception: *r* = −0.61, *p* < 0.001], suggesting a greater reliance on online adjustments and ongoing decision-making after movement initiation (Fig. 4C). Furthermore, there were RT differences depending on the movement strategy (redirect-to-avoid, direct avoid, direct object hit) ultimately executed. Redirect-to-avoid movements (i.e., movements initiated towards ellipse but subsequently corrected) had an average RT of 390 ± 11 ms, relative to 489 ± 15 ms for direct avoids [main effect of movement strategy: *F*(1.32, 33.12) = 71.64, *p* < 0.001, η^2^ = 0.35, Greenhouse-Geisser corrected] (Fig. 4D). The average RT for direct object hits was approximately halfway in-between the RTs for the two types of avoid movements (439 ± 12 ms), reflecting that participants defaulted towards initiating a movement towards the object even when their decision was incomplete. Interestingly, RTs were shorter for Interception than Reaching for redirect-to-avoid, direct avoids, and direct hits [all *t*’s > 2.2, all *p*’s < 0.05], and the RT difference was largest for direct avoids [interaction of movement type and strategy: *F*(1.34, 33.48) = 8.51, *p* = 0.003, η^2^ = 0.02, Greenhouse-Geisser corrected]. This suggests that simply preparing for an interception movement, even when it was not selected, contributed to earlier movement initiation.

A closer analysis of the movement trajectories suggests that the initial movement plans carried a signature of an incomplete decision during movement initiation. Both direct object hit and redirect-to-avoid movements were initially aimed toward the object, indicating an early motor plan to hit the object. However, as shown in Figure 4E, trajectories of movements that were ultimately redirected were on average initially aimed farther from the midline than direct movements (longer tail for redirect-to-avoid) [main effect of movement strategy: *F*(1,25) = 131.91, *p* < 0.001, η^2^ = 0.28], and this difference was larger for Interception [interaction of movement type and strategy: *F*(1, 25) = 10.59, *p* = 0.003, η^2^ = 0.01]. The deviation of the initial direction away from the midline likely reflects an intermediate motor plan between hitting the circle and the bar, suggesting a more conservative approach when the decision is not fully formed.

### Perceptual decision-making influences eye movement strategies

Saccades and gaze events were identified using a geometric method to transform eye movement data to the horizontal plane and adaptive velocity-based thresholds (Singh et al. 2016) for each participant (see Fig. 5A). Standard task performance consisted of an initial saccade followed by onset of gaze (fixation or smooth-pursuit) on the target - we restricted our eye movement analysis to the trials that followed that structure (see Methods for details).

**Figure 5:**
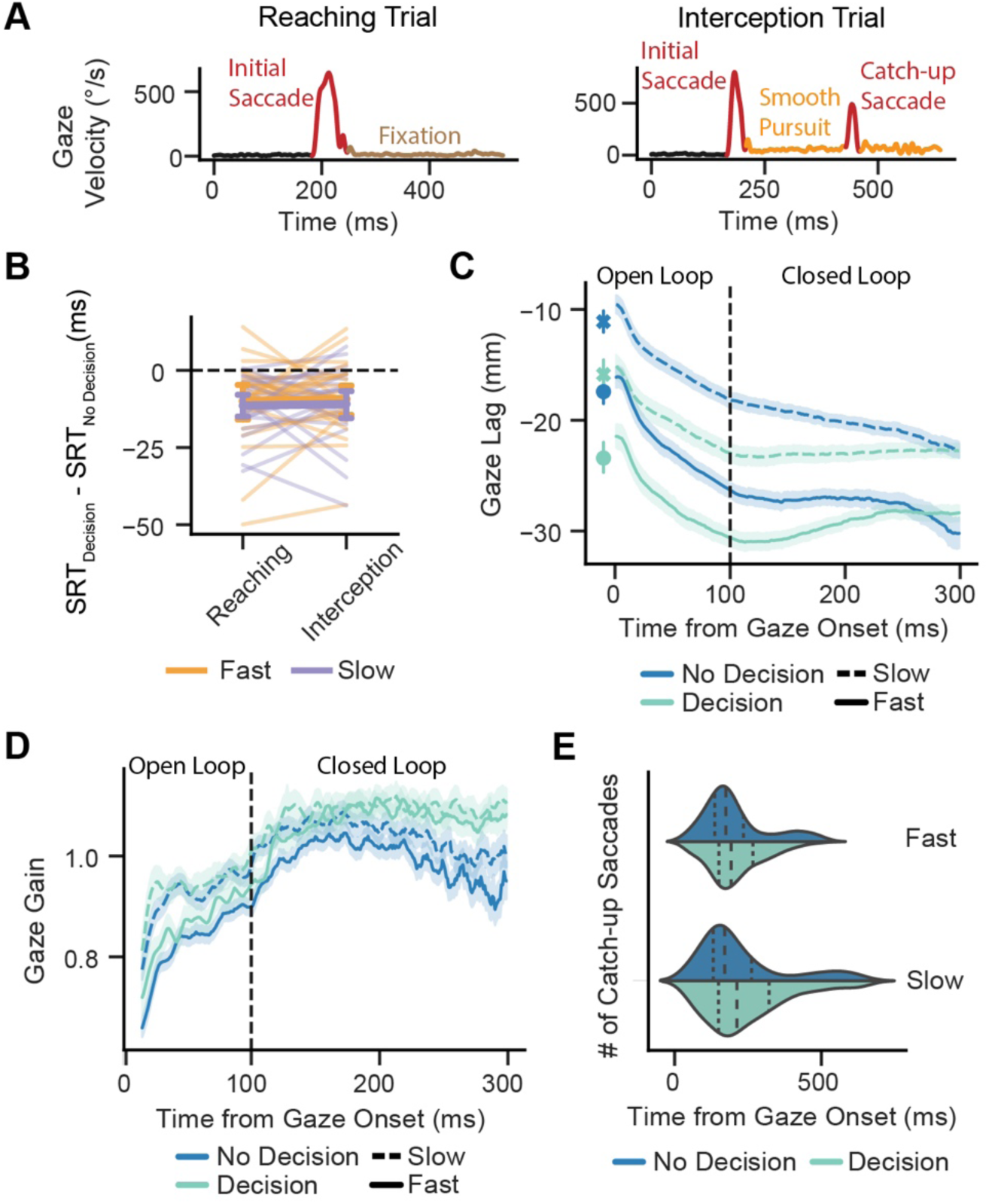
Perceptual decision-making influences eye movement strategies. A: Two representative trials showing classification of gaze events using adaptive velocity-based thresholds for reaching to stationary targets and intercepting moving targets. B: Saccadic reaction times decreased for Decision blocks similarly for reaching and interception. C: Gaze lag across interception trials for the end of the initial saccade and as a function of time from gaze onset. Positive values indicate that the gaze led the object, whereas negative values indicate lag. Participants lagged more during Decision blocks and at fast trial durations (higher object velocity). The error bars and shaded area represents the 95% confidence interval of the mean estimate. D: Gaze gain across interception trials as a function of time from gaze onset. Gaze gain was higher for faster velocities during the open-loop period and higher for Decision blocks during the closed-loop period. The shaded area represents the 95% confidence interval of the mean estimate. E: Distribution of catch-up saccades after gaze onset during interception. Catch-up saccades were more frequent during Decision blocks. The dotted lines denote the quartiles of the distribution.

As shown in Figure 5B, SRTs during Decision blocks were on average 10.4 (± 2.3) ms faster during Decision blocks than No Decision blocks [*t*(1,23) = −5.52, *p <* 0.001]. The decrease in SRTs was similar for both Reaching and Interception [main effect of movement type: *F*(1,23) = 0.08 *p* = 0.79, η^2^ < 0.01], suggesting that adding a perceptual decision increased the general urgency to launch a saccade. However, as can be seen for Interception movements, there was likely a speed-accuracy trade-off associated with faster SRTs: the initial saccade landed farther behind the moving object during Decision blocks [see Fig. 5C; main effect of decision: *F*(1,23) = 13.93, *p* = 0.001, η^2^ = 0.04] and for Fast trial durations (i.e., when the object was moving at faster velocities) [main effect of trial duration: *F*(1,23) = 37.25, *p* < 0.001, η^2^ = 0.07]. Eye position lag persisted during approximately the first 300 ms of the smooth-pursuit period [main effect of decision type: *F*(1,23) = 17.74, *p* < 0.001, η^2^ = 0.14; main effect of trial duration: *F*(1,23) = 114.18, *p* < 0.001, η^2^ = 0.32]. This result suggests that the urgency of the initial saccade led to less precise oculomotor movement during decision-making.

Participants compensated for the initial lag in pursuit by increasing the gaze gain. Though gaze gain in the open-loop period (15-100 ms after pursuit initiation) was driven mainly by differences in object velocity [main effect of trial duration: *F*(1,23) = 58.67, *p* < 0.001, η^2^ = 0.13], during the closed-loop period gaze gain increased for Decision blocks relative to No Decision blocks [main effect of decision: *F*(1,23) = 49.02, *p* < 0.001, η^2^ = 0.15] (Fig. 5D). This effect is not simply due to longer pursuit durations during Decision blocks, as gains are also longer when the analysis is restricted to the first 100 ms of the closed-loop period [main effect of decision: *F*(1,23) = 8.71, *p* = 0.007, η^2^ = 0.04]. This suggests that the negative closed feedback loop that minimizes retinal error between gaze and target is engaged differently when perceptual decision-making task-constraints are imposed during pursuit eye movements.

Participants also initiated more catch-up saccades during Decision blocks (*M =* 0.99 ± 0.30 saccades/s) than No Decision blocks (*M =* 0.68 ± 0.30 saccades/s) to make up for the lag in object pursuit [main effect of decision type: *F*(1,23) = 16.23, *p* < 0.001, η^2^ = 0.10] (see Fig. 5E). The mean latency of when the catch-up saccade occurred relative to pursuit onset did not differ across decision type blocks [main effect of decision: *F*(1,23) = 1.06, *p* = 0.31, η^2^ < 0.01] or trial duration [main effect of trial duration: *F*(1,23) = 2.25, *p* = 0.15, η^2^ = 0.02]. Together, these results suggest that ocular movements are altered when decision about object features have to be made in addition to estimating its spatial location.

## Discussion

In the current study, we asked the question: how does perceptual decision-making involving the two visual streams affect visuomotor coordination during reaching and interception movements? To address this question, we manipulated ventral stream involvement in a rapid visuomotor task. In one condition, participants made reaching or interception movements to hit an object shaped like a circle. In another condition, participants had to judge the shape of the object: if a circle appeared, they were instructed to reach or intercept it, but if an ellipse appeared, they were instead instructed to make a movement away from the ellipse and towards a horizontal bar. Our results support our first hypothesis of differential effects of ventral stream engagement on dorsal stream processing during interception relative to reaching movements. Furthermore, we also found support for our second hypothesis - that changes in oculomotor behavior when the ventral visual stream is engaged may contribute to differences in limb motor performance.

Many studies have probed the interactions between dorsal and ventral stream processes during reaching movements (reviewed in Song and Nakayama 2009) but to the best of our knowledge only a handful of studies have extended this type of paradigm to interception movements (de la Malla et al. 2019; Lacquaniti and Maioli 1989). Our approach also differs from the classical *backward masking* approach used by some researchers to quantify how object recognition affects planning and execution of reaching movements (Cressman et al. 2007; Schmidt 2002). In this approach, a brief target stimulus (prime) is followed by a mask that impedes recognition of the target. These studies showed that reaching movement trajectories were strongly affected by the prime target, even when blocked from awareness by masking, suggesting a flow of object property information from the ventral visual stream to the dorsal action stream. In our approach, we presented the same stimulus for the entire trial duration to afford participants flexibility in how they processed object shape. We chose two trial times of 800 ms (Fast) and 950 ms (Slow) to give participants enough time to identify object shape (∼250-300 ms) and plan movements (∼100-200 ms) in a sequential fashion, i.e. to minimize decision errors participants could first ascertain the object shape and then plan the movement trajectory. Our paradigm also allowed participants to judge the object shape and prepare a motor plan simultaneously. If the slower ventral stream process of shape recognition took longer than the preparation of the motor plan, we predicted that effective ventral-dorsal stream integration would allow participants to take corrective action by completing shape recognition after the movement had been initiated. Our results show that participants used both strategies. Longer reaction times of ∼500 ms were associated with fewer decision errors and redirected movements (see Fig. 3C and 4C). In contrast, average reaction times of ∼400 ms were associated with more decision errors as well as corrective redirected movements.

### Online integration of ventral stream and decision processing during interception

Vision for goal selection based on object properties and vision guiding the online control of movement have been conceptualized as two specialized processes mediated by the ventral and dorsal streams, respectively (Goodale and Milner 1992; Goodale and Westwood 2004). While much work has concerned how the two visual streams serve unique functional roles operating largely independent of each other, less is known about the interaction in more complex task environments. The current task was designed to force this interaction—that is, in order to perform the correct action (hit the object or avoid it), participants must accurately identify the object’s shape (circle or ellipse). We found that even under time constraints (800 ms to hit the object in the Fast condition), participants could recognize objects and formulate a decision prior to movement initiation. Relative to No Decision blocks, in which participants only needed to process spatial information to facilitate movement, there was an average RT delay of 178 ms in Decision blocks (see Fig. 3B), suggesting additional processing time for shape recognition and motor goal selection (Cisek and Kalaska 2010; Thorpe and Fabre-Thorpe 2001; Veerman et al. 2008). Thus, it is reasonable to assume from the average RTs that perceptual processing in the ventral stream could precede dorsal stream processing of visuomotor transformations for action execution.

However, closer investigation of the movement trajectories and corresponding RTs provides evidence that processing of object information and decision-making continues after movement initiation. During both reaching and interception, we observed that participants would often initiate their movements toward the circle only to curve around past the original start location and hit the bar. The presence of these “redirect-to-avoid” movements (see Fig. 4D) provide evidence of an evolving decision given accumulating stimulus information (Resulaj et al. 2009; Selen et al. 2012). In contrast to previous studies investigating sensorimotor decisions of the limb that vary the motion or spatial location of the target (Burk et al. 2014; Gallivan et al. 2016; van den Berg et al. 2016), here we show that sensorimotor transformations computed in the dorsal stream can seamlessly integrate incoming information about object shape that originates in the ventral stream (Davare et al. 2007; Konen and Kastner 2008; Lehky and Tanaka 2016; Sereno and Maunsell 1998). The distribution of initial movement directions (see Fig. 4E) of redirected movements toward the direction of the bar suggests that movements are planned to optimize task success given uncertainty about the impending decision (Haith et al. 2015; Nashed et al. 2017; Wong and Haith 2017). Thus, even though the imposed time constraints allowed for sequential stimulus identification, decision-making, and movement execution, participants tended to favor an alternative strategy in which both these processes co-occurred during preparation and execution (Haith et al. 2016; Orban de Xivry et al. 2017).

What determines the reliance on integration of ventral and dorsal stream information during visuomotor control? In the present task, the complexity of the motor response modulated the perceived urgency to act (Thura 2020; Thura and Cisek 2016). Both initial and final decision errors increased during interception relative to reaching during decision-making, largely due to participants initially aiming toward and then unable to correct a response toward a moving ellipse. In addition, movements were more likely to be redirected during interception, indicating a stronger bias toward initiating a hit movement prior to making a perceptual decision about object shape. Furthermore, an individual’s initial decision error rate and tendency to perform redirect movements were each associated with shorter RTs, indicating that the shorter RTs during interceptions in Decision blocks were likely due to a greater dependency on online decision-making and motor control (Brenner and Smeets 2018).

However, given that the urgency to act during interception had clear consequences on task performance (more decisional errors), the capacity for integration of ventral stream information with visuomotor performance may be limited. Our results suggest that the urgency of the response may interfere with, rather than be a consequence of, differential ventral-dorsal stream interactions. Further work directly addressing different stimulus attributes associated with separate areas along the ventral pathway (e.g., orientation, color, size) can help clarify how movements are planned relative to the time-course of sensory processing and decision-making. Notably, the errors in interception during decision-making were associated with the inability to adjust initial movement trajectories that account for decisional demands, but the increase in aiming errors was no different between interception and reaching. This suggests that the interference in the time-course of ventral-dorsal stream interactions mainly affects decision processes rather than the online control of movement per se.

Our study does not address how the dorsal stream receives ventral stream information about object shape, but recent work has identified pathways between the two streams that could facilitate direct communication during ongoing sensorimotor control (Budisavljevic et al. 2018; Takemura et al. 2016). The present findings suggest that that the motor system can integrate prolonged processing of sensory information originating in the ventral stream, but how the extent to which this integrated information can be accessed depends on movement complexity.

### Modulation of gaze gains during perceptual decision-making

During Decision blocks, saccades were launched about 10 ms earlier than No-Decision blocks. It appears that the earlier launch of the saccade was because of a perceived urgency to recognize the object shape and make the correct motor decision. Saccades to visible targets are generally imprecise and undershoot target position (Krappmann 1998). Thus, the earlier launch may have occurred before the spatial planning of the saccade was complete, resulting in larger undershoots farther away from the object (larger gaze lags in Decision blocks, Fig. 5C). Since in our study objects had to be foveated to be recognized, the oculomotor system may have increased the gaze gains (Fig. 5D) and made more catch-up saccades (Fig. 5E) to the target during Decision blocks to compensate for the large lags at the end of the saccades.

Smooth-pursuit gains have been conventionally defined as the ratio of target and gaze velocity in angular coordinates in head-fixed conditions. The first 100 ms of the smooth-pursuit movement is referred to as the open-loop phase (Barnes 2008; Tychsen and Lisberger 1986). This is followed by the onset of closed-loop pursuit, which is mainly controlled by a negative feedback loop to ensure that the eye velocity closely matches the target velocity. However, pursuit gains are defined for head-fixed conditions to ensure that the vestibular-ocular reflex does not interfere with gaze movements. Since our eye-tracker could have allowed small head movements, we decided to report gaze gains (Barnes 1993; Collins and Barnes 1999; Ranalli and Sharpe 1988) instead of pursuit gains. One study in primates has shown that when the head is unrestrained, pursuit and gaze gains are similar suggesting that eye and head movements are controlled together within the pursuit pathways (Dubrovsky and Cullen 2002). Thus, we compared both open-loop (first 100 ms) and closed-loop gaze gains (>100 ms) as a proxy for pursuit gains for the Interception blocks for the No Decision and Decision conditions.

As expected, changes in the open-loop gains were driven predominantly by object velocity (Fast versus Slow). However, the closed-loop gains were significantly higher for the Decision than No Decision blocks. An important question is whether these higher gains for the Decision blocks reflected the constraints imposed by shape recognition or were simply a compensation for the large errors in where the saccade landed. Previously, it has been shown that object recognition is impaired when targets move at high speeds (Ludvigh and Miller 1958b; Schütz et al. 2009; Westheimer and McKee 1975). In contrast to the slow speed of 1-10°/sec used in these studies, the objects in our experiment moved at approximately 80-90°/sec. This speed approaches the limit of smooth-pursuit in humans (Meyer et al. 1985) and we expected that participants would not only have trouble in pursuing objects at high speeds, but that it would also compromise their ability to recognize objects. However, the closed-loop pursuit gains were similar between Fast and Slow blocks, and only differed between the Decision blocks. Thus, it seems that the gaze lag (caused by earlier release of the saccade) and the need to foveate the object to recognize the shape together contributed to a higher closed-loop gaze gain. This suggests that the negative closed feedback loop that minimizes retinal error between gaze and target is engaged differently when the ventral stream is engaged for perceptual decision-making during pursuit eye movements.

Our result suggests that the visual perceptual decision-making network, that includes the ventral visual stream, dorsolateral prefrontal regions and frontal eye fields (Heekeren et al. 2004; Heekeren et al. 2008; Sakagami and Pan 2007), may provide either a predictive or urgency signal to the smooth-pursuit system to increase the gain and minimize the retinal error between the target and the gaze. Indeed, stimulation and lesion studies have implicated the frontal eye fields with the modulation of smooth-pursuit gain during object tracking (Gagnon et al. 2006; Keating 1991; Morrow and Sharpe 1995; Shi et al. 1998). Furthermore, anatomical tracer studies in primates have shown that the dorsal and ventral processing streams converge in the lateral frontal eye fields (Schall et al. 1995). Taken together with our data, this suggests that in tasks where perceptual decision-making is necessary during pursuit eye movements, the frontal eye fields may modulate gaze gains to meet task demands.

## Conclusions

In this study, we introduced a visuomotor decision-making task in which a successful reaching or interception movement depended on visual processing for perception and action in the ventral and dorsal streams. We found that engagement of the ventral stream led to more decision errors and a smaller increase in hand RTs for interception movements relative to reaching movements, reflective of a greater perceived urgency to act during interception. During decision-making, participants had faster saccadic RTs and adopted online movement strategies that incorporated an evolving decision about object shape. Additionally, participants exhibited higher gaze gains to adapt to the demands of integrating the perceptual decision with visuomotor control. These results suggest that the capacity to effectively integrate ventral-dorsal stream information during ongoing movement depends on the perceived urgency to act, which is greater when intercepting a moving target.

## Acknowledgements

We thank Negar Bassiri and Chris Mejias for assisting with data collection. DAB received support from the American Heart Association (18POST34060183). A portion of this research was supported by a grant from the University of Georgia Research Foundation, Inc to TS.

## Notes

**Disclosures:** The authors declare no conflict of interest, financial or otherwise.

#### Summary of Updates

Article updated after comments from two Reviewers.

## References

Ales JM, Appelbaum LG, Cottereau BR, and Norcia AM. The time course of shape discrimination in the human brain. Neuroimage 67: 77–88, 2013.

Barnes GR. Cognitive processes involved in smooth pursuit eye movements. Brain and Cognition 68: 309–326, 2008.

Barnes GR. Visual-vestibular interaction in the control of head and eye movement: The role of visual feedback and predictive mechanisms. Progress in Neurobiology 41: 435–472, 1993.

Brenner E, and Smeets JB. Sources of variability in interceptive movements. Exp Brain Res 195: 117–133, 2009.

Brenner E, and Smeets JBJ. Continuously updating one’s predictions underlies successful interception. J Neurophysiol 120: 3257–3274, 2018.

Brostek L, Eggert T, and Glasauer S. Gain Control in Predictive Smooth Pursuit Eye Movements: Evidence for an Acceleration-Based Predictive Mechanism. eNeuro 4: 2017.

Budisavljevic S, Dell’Acqua F, and Castiello U. Cross-talk connections underlying dorsal and ventral stream integration during hand actions. Cortex 103: 224–239, 2018.

Burk D, Ingram JN, Franklin DW, Shadlen MN, and Wolpert DM. Motor effort alters changes of mind in sensorimotor decision making. PLoS One 9: e92681, 2014.

Churchland AK, and Lisberger SG. Gain control in human smooth-pursuit eye movements. J Neurophysiol 87: 2936–2945, 2002.

Cisek P, and Kalaska JF. Neural mechanisms for interacting with a world full of action choices. Annu Rev Neurosci 33: 269–298, 2010.

Collins C, and Barnes G. Independent control of head and gaze movements during head-free pursuit in humans. Journal of Physiology 515: 299–314, 1999.

Cressman EK, Franks IM, Enns JT, and Chua R. On-line control of pointing is modified by unseen visual shapes. Consciousness and Cognition 16: 265–275, 2007.

Culham JC, Cavina-Pratesi C, and Singhal A. The role of parietal cortex in visuomotor control: what have we learned from neuroimaging? Neuropsychologia 44: 2668–2684, 2006.

Davare M, Andres M, Clerget E, Thonnard JL, and Olivier E. Temporal dissociation between hand shaping and grip force scaling in the anterior intraparietal area. J Neurosci 27: 3974–3980, 2007.

Day BL, and Lyon IN. Voluntary modification of automatic arm movements evoked by motion of a visual target. Exp Brain Res 130: 159–168, 2000.

de la Malla C, Brenner E, de Haan EHF, and Smeets JBJ. A visual illusion that influences perception and action through the dorsal pathway. Communications Biology 2: 38, 2019.

Diedenhofen B, and Musch J. cocor: a comprehensive solution for the statistical comparison of correlations. PLoS One 10: e0121945, 2015.

Dubrovsky AS, and Cullen KE. Gaze-, eye-, and head-movement dynamics during closed-and open-loop gaze pursuit. Journal of Neurophysiology 87: 859–875, 2002.

Dunn OJ, and Clark V. Correlation coefficients measured on the same individuals. Journal of the American Statistical Association 64: 366–377, 1969.

Fooken J, and Spering M. Decoding go/no-go decisions from eye movements. J Vis 19: 5, 2019.

Franklin DW, Reichenbach A, Franklin S, and Diedrichsen J. Temporal Evolution of Spatial Computations for Visuomotor Control. J Neurosci 36: 2329–2341, 2016.

Gagnon D, Paus T, Grosbras M-H, Pike GB, and O’Driscoll GA. Transcranial magnetic stimulation of frontal oculomotor regions during smooth pursuit. Journal of Neuroscience 26: 458–466, 2006.

Gallivan JP, Chapman CS, Wolpert DM, and Flanagan JR. Decision-making in sensorimotor control. Nat Rev Neurosci 19: 519–534, 2018.

Gallivan JP, and Goodale MA. The dorsal “action” pathway. In: Handbook of clinical neurologyElsevier, 2018, p. 449–466.

Gallivan JP, Logan L, Wolpert DM, and Flanagan JR. Parallel specification of competing sensorimotor control policies for alternative action options. Nat Neurosci 19: 320–326, 2016.

Gauthier GM, Nommay D, and Vercher JL. The role of ocular muscle proprioception in visual localization of targets. Science 249: 58–61, 1990.

Gold JI, and Shadlen MN. The neural basis of decision making. Annu Rev Neurosci 30: 535–574, 2007.

Goodale MA, and Milner AD. Separate visual pathways for perception and action. Trends Neurosci 15: 20–25, 1992.

Goodale MA, and Westwood DA. An evolving view of duplex vision: separate but interacting cortical pathways for perception and action. Curr Opin Neurobiol 14: 203–211, 2004.

Grill-Spector K, Kourtzi Z, and Kanwisher N. The lateral occipital complex and its role in object recognition. Vision Res 41: 1409–1422, 2001.

Gritsenko V, Yakovenko S, and Kalaska JF. Integration of predictive feedforward and sensory feedback signals for online control of visually guided movement. J Neurophysiol 102: 914–930, 2009.

Haith AM, Huberdeau DM, and Krakauer JW. Hedging your bets: intermediate movements as optimal behavior in the context of an incomplete decision. PLoS Comput Biol 11: e1004171, 2015.

Haith AM, Pakpoor J, and Krakauer JW. Independence of Movement Preparation and Movement Initiation. J Neurosci 36: 3007–3015, 2016.

Hecht D, Reiner M, and Karni A. Multisensory enhancement: gains in choice and in simple response times. Exp Brain Res 189: 133–143, 2008.

Heekeren HR, Marrett S, Bandettini PA, and Ungerleider LG. A general mechanism for perceptual decision-making in the human brain. Nature 431: 859–862, 2004.

Heekeren HR, Marrett S, and Ungerleider LG. The neural systems that mediate human perceptual decision making. Nature Reviews Neuroscience 9: 467–479, 2008.

Holm S. A Simple Sequentially Rejective Multiple Test Procedure. Scandinavian Journal of Statistics 6: 65–70, 1979.

Joo SJ, Katz LN, and Huk AC. Decision-related perturbations of decision-irrelevant eye movements. Proc Natl Acad Sci U S A 113: 1925–1930, 2016.

Keating E. Frontal eye field lesions impair predictive and visually-guided pursuit eye movements. Experimental Brain Research 86: 311–323, 1991.

Konen CS, and Kastner S. Two hierarchically organized neural systems for object information in human visual cortex. Nat Neurosci 11: 224–231, 2008.

Krappmann P. Accuracy of visually and memory-guided antisaccades in man. Vision Research 38: 2979–2985, 1998.

Lacquaniti F, and Maioli C. The role of preparation in tuning anticipatory and reflex responses during catching. Journal of Neuroscience 9: 134–148, 1989.

Lehky SR, and Tanaka K. Neural representation for object recognition in inferotemporal cortex. Curr Opin Neurobiol 37: 23–35, 2016.

Lencer R, and Trillenberg P. Neurophysiology and neuroanatomy of smooth pursuit in humans. Brain Cogn 68: 219–228, 2008.

Lisberger SG. Visual Guidance of Smooth Pursuit Eye Movements. Annu Rev Vis Sci 1: 447–468, 2015.

Ludvigh E, and Miller JW. Study of visual acuity during the ocular pursuit of moving test objects. I. Introduction. J Opt Soc Am 48: 799–802, 1958a.

Ludvigh E, and Miller JW. Study of visual acuity during the ocular pursuit of moving test objects. I. Introduction. Journal of the Optical Society of America 48: 799–802, 1958b.

Merchant H, Zarco W, Prado L, and Perez O. Behavioral and neurophysiological aspects of target interception. In: Progress in Motor Control Advances in Experimental Medicine and Biology, edited by Sternad D. Boston, MA: Springer, 2009, p. 201–220.

Meyer CH, Lasker AG, and Robinson DA. The upper limit of human smooth pursuit velocity. Vision Research 25: 561–563, 1985.

Milner AD. How do the two visual streams interact with each other? Experimental brain research 235: 1297–1308, 2017.

Mishkin M, Ungerleider LG, and Macko KA. Object vision and spatial vision: two cortical pathways. Trends in Neurosciences 6: 414–417, 1983.

Morrow MJ, and Sharpe JA. Deficits of smooth-pursuit eye movement after unilateral frontal lobe lesions. Annals of Neurology 37: 443–451, 1995.

Nashed JY, Diamond JS, Gallivan JP, Wolpert DM, and Flanagan JR. Grip force when reaching with target uncertainty provides evidence for motor optimization over averaging. Sci Rep 7: 11703, 2017.

Orban de Xivry JJ, Legrain V, and Lefevre P. Overlap of movement planning and movement execution reduces reaction time. J Neurophysiol 117: 117–122, 2017.

Ranalli PJ, and Sharpe JA. Vertical vestibulo-ocular reflex, smooth pursuit and eye-head tracking dysfunction in internuclear ophthalmoplegia. Brain 111: 1299–1317, 1988.

Resulaj A, Kiani R, Wolpert DM, and Shadlen MN. Changes of mind in decision-making. Nature 461: 263–266, 2009.

Rizzolatti G, Fogassi L, and Gallese V. Motor and cognitive functions of the ventral premotor cortex. Curr Opin Neurobiol 12: 149–154, 2002.

Rizzolatti G, and Matelli M. Two different streams form the dorsal visual system: anatomy and functions. Exp Brain Res 153: 146–157, 2003.

Rosenbaum DA, Cohen RG, Jax SA, Weiss DJ, and Van Der Wel R. The problem of serial order in behavior: Lashley’s legacy. Human Movement Science 26: 525–554, 2007.

Sakagami M, and Pan X. Functional role of the ventrolateral prefrontal cortex in decision making. Current Opinion in Neurobiology 17: 228–233, 2007.

Sarlegna FR, and Mutha PK. The influence of visual target information on the online control of movements. Vision Res 110: 144–154, 2015.

Schall JD, Morel A, King DJ, and Bullier J. Topography of visual cortex connections with frontal eye field in macaque: convergence and segregation of processing streams. Journal of Neuroscience 15: 4464–4487, 1995.

Schmidt T. The finger in flight: real-time motor control by visually masked color stimuli. Psychol Sci 13: 112–118, 2002.

Schutz AC, Braun DI, and Gegenfurtner KR. Object recognition during foveating eye movements. Vision Research 49: 2241–2253, 2009.

Schütz AC, Braun DI, and Gegenfurtner KR. Object recognition during foveating eye movements. Vision Research 49: 2241–2253, 2009.

Schwartz EL, Desimone R, Albright TD, and Gross CG. Shape recognition and inferior temporal neurons. Proc Natl Acad Sci U S A 80: 5776–5778, 1983.

Selen LP, Shadlen MN, and Wolpert DM. Deliberation in the motor system: reflex gains track evolving evidence leading to a decision. J Neurosci 32: 2276–2286, 2012.

Sereno AB, and Maunsell JH. Shape selectivity in primate lateral intraparietal cortex. Nature 395: 500–503, 1998.

Shi D, Friedman HR, and Bruce CJ. Deficits in smooth-pursuit eye movements after muscimol inactivation within the primate’s frontal eye field. Journal of Neurophysiology 80: 458–464, 1998.

Singh T, Fridriksson J, Perry CM, Tryon SC, Ross A, Fritz S, and Herter TM. A novel computational model to probe visual search deficits during motor performance. J Neurophysiol 117: 79–92, 2017.

Singh T, Perry CM, and Herter TM. A geometric method for computing ocular kinematics and classifying gaze events using monocular remote eye tracking in a robotic environment. Journal of Neuroengineering and Rehabilitation 13: 10, 2016.

Song JH, and Nakayama K. Hidden cognitive states revealed in choice reaching tasks. Trends Cogn Sci 13: 360–366, 2009.

Song JH, and Nakayama K. Target selection in visual search as revealed by movement trajectories. Vision Research 48: 853–861, 2008.

Spering M, Kerzel D, Braun DI, Hawken MJ, and Gegenfurtner KR. Effects of contrast on smooth pursuit eye movements. J Vis 5: 455–465, 2005.

Takemura H, Rokem A, Winawer J, Yeatman JD, Wandell BA, and Pestilli F. A Major Human White Matter Pathway Between Dorsal and Ventral Visual Cortex. Cereb Cortex 26: 2205–2214, 2016.

Thorpe SJ, and Fabre-Thorpe M. Neuroscience. Seeking categories in the brain. Science 291: 260–263, 2001.

Thura D. Decision urgency invigorates movement in humans. Behav Brain Res 382: 112477, 2020.

Thura D, and Cisek P. Modulation of Premotor and Primary Motor Cortical Activity during Volitional Adjustments of Speed-Accuracy Trade-Offs. J Neurosci 36: 938–956, 2016.

Tychsen L, and Lisberger SG. Visual motion processing for the initiation of smooth-pursuit eye movements in humans. Journal of Neurophysiology 56: 953–968, 1986.

van den Berg R, Anandalingam K, Zylberberg A, Kiani R, Shadlen MN, and Wolpert DM. A common mechanism underlies changes of mind about decisions and confidence. Elife 5: e12192, 2016.

van Polanen V, and Davare M. Interactions between dorsal and ventral streams for controlling skilled grasp. Neuropsychologia 79, Part B: 186–191, 2015.

Veerman MM, Brenner E, and Smeets JB. The latency for correcting a movement depends on the visual attribute that defines the target. Exp Brain Res 187: 219–228, 2008.

Westheimer G, and McKee SP. Visual acuity in the presence of retinal-image motion. Journal of the Optical Society of America 65: 847–850, 1975.

Wong AL, and Haith AM. Motor planning flexibly optimizes performance under uncertainty about task goals. Nature Communications 8: 14624, 2017.

Zago M, McIntyre J, Senot P, and Lacquaniti F. Visuo-motor coordination and internal models for object interception. Exp Brain Res 192: 571–604, 2009.

